# Draft Genomes of two *Artocarpus* plants, Jackfruit (*A. heterophyllus*) and Breadfruit (*A. altilis*)

**DOI:** 10.1101/869339

**Authors:** Sunil Kumar Sahu, Min Liu, Anna Yssel, Robert Kariba, Sanjie Jiang, Bo Song, Samuel Muthemba, Prasad S. Hendre, Ramni Jamnadass, Shu-Min Kao, Jonathan Featherston, Nyree J.C. Zerega, Xun Xu, Huanming Yang, Allen Van Deynze, Yves Van de Peer, Xin Liu, Huan Liu

**Affiliations:** BGI-Shenzhen, Shenzhen 518083, China; State Key Laboratory of Agricultural Genomics, BGI-Shenzhen, Shenzhen 518083, China; Center for Microbial Ecology and Genomics (CMEG); Department of Biochemistry, Genetics and Microbiology; University of Pretoria, Pretoria, South Africa; African Orphan Crops Consortium, World Agroforestry Centre (ICRAF), Nairobi, Kenya; Department of Plant Biotechnology and Bioinformatics, Ghent University, Belgium; Center for Plant Systems Biology, VIB, Belgium; Biotechnology Platform, Agricultural Research Council, Pretoria, 0110, South Africa; Chicago Botanic Garden, Negaunee Institute for Plant Conservation Science and Action, Glencoe, IL, 60022, USA; Northwestern University, Plant Biology and Conservation, Evanston, IL, 60208, USA; University of California, 1 Shields Ave, Davis, USA, 95616; Department of Biology, University of Copenhagen, Copenhagen, Denmark

**Author notes:** Equal contribution (Sunil Kumar Sahu, Min Liu). Correspondence: Huan Liu; Xin Liu.

**Keywords:** Jackfruit, Breadfruit, *A. heterophyllus*, *A. altilis*, starch synthesis

## Abstract

Two of the most economically important plants in the *Artocarpus* genus are jackfruit (*A. heterophyllus* Lam.) and breadfruit (*A. altilis* (Parkinson) Fosberg). Both species are long-lived trees that have been cultivated for thousands of years in their native regions. Today they are grown throughout tropical to subtropical areas as an important source of starch and other valuable nutrients. There are hundreds of breadfruit varieties that are native to Oceania, of which the most commonly distributed types are seedless triploids. Jackfruit is likely native to the western Ghats of India and produces one of the largest tree-borne fruit structures (reaching up to 100 pounds). To date, there is limited genomic information for these two economically important species. Here, we generated 273 Gb and 227 Gb of raw data from jackfruit and breadfruit, respectively. The high-quality reads from jackfruit were assembled into 162,440 scaffolds totaling 982 Mb with 35,858 genes. Similarly, the breadfruit reads were assembled into 180,971 scaffolds totaling 833 Mb with 34,010 genes. A total of 2,822 and 2,034 expanded gene families were found in jackfruit and breadfruit, respectively, enriched in pathways including starch- and sucrose metabolism, photosynthesis and others. The copy number of several starch synthesis related genes were found increased in jackfruit and breadfruit compared to closely related species, and the tissue specific expression might imply their sugar-rich and starch-rich characteristics. Overall, the publication of high-quality genomes for jackfruit and breadfruit provides information about their specific composition and the underlying genes involved in sugar and starch metabolism.

## 1. Introduction

The family Moraceae contains at least 39 genera and approximately 1,100 species [1–3]. Species diversity of the family is primarily centered around the tropics with variation in inflorescence structures, pollination forms, breeding systems and growth forms [2]. Within the Moraceae family, the genus *Artocarpus* is comprised of approximately 70 species [2,4]. The most recent evidence indicates that Borneo was the center of diversification of the *Artocarpus* genus and that species diversified throughout South and Southeast Asia [2]. All members of the genus have unisexual flowers and produce exudate from laticifers. Inflorescences consist of up to thousands of tiny flowers, tightly packed and condensed on a receptacle [2]. In most species, the perianths of adjacent female flowers are partially to completely fused together and develop into a highly specialized multiple fruit called a syncarp, which is formed by the enlargement of the entire female head. Syncarps of different species range in size from a few centimeters in diameter to over half a meter long in the case of jackfruit. [2,5]. Many *Artocarpus* species are important food sources for forest fauna, and about a dozen species are important crops in the regions where they are from [2,6].

Jackfruit (*A. heterophyllus* Lam.) grows wild in the Western Ghats of India and is cultivated as an important food source across the tropics. It is monecious, and thought to be pollinated by gall midges 7]. In some areas it is propagated mainly by seeds [8], however, clonal propagation via grafting is increasing in areas where it is grown for commercial use [9]. On an average it contains more than 100 seeds per fruit with viability of less than a month [10,11]. The male flowers are tiny and clustered on an oblong receptacle, typically 2 to 4 inches (5-10 cm) in length. Limited studies exist on the range of cultivated varieties of jackfruit, but they are often grouped into two main types, varieties with edible fleshy perianth tissue (often referred to as “flakes”) that are either (a) small, fibrous, soft and spongy or (b) larger, less sweet fruit with crisp [10,12]. The latter type is often more commercially important.

Breadfruit [*A. altilis* (Parkinson) Fosberg] is most likely derived from the progenitor species *A. camansi* Blanco, which is native to New Guinea [10,13]. As humans migrated and colonized the islands of Remote Oceania, indigenous people selected and cultivated varieties from the wild ancestor over thousands of years [13], giving rise to hundreds of cultivated varieties [10,13–15]. Cultivated varieties were traditionally propagated clonally by root cuttings but can now be commercially propagated by tissue culture [16,17]. Among the hundreds of varieties, some are diploid (2n=2x=~56) and may produce seeds, while other varieties are seedless triploids (3n=2x=~84), and still others are of hybrid origin with another species, *A. mariannensis* Trécul [13,18–20]. A small subset of the triploid diversity is what has been introduced outside of Oceania [19,21].

To diversify the global food supply, enhance agricultural productivity and tackle malnutrition, it is necessary focus more on crop plants that are utilized in rural societies as a local source of nutrition and sustenance, but have received little attention for crop improvement. This study is part of the African Orphan Crops Consortium (AOCC), an international public-private partnership. A goal of this global initiative is to sequence, assemble and annotate the genomes of 101 traditional African food crops by 2020 [22,23]. Both breadfruit and jackfruit are nutritious [24–27] and have potential to increase food security, especially in tropical areas. Until now only limited genomic information has been available for the *Artocarpus* genus as a whole. Microsatellite markers have been used to characterize cultivars and wild relatives of breadfruit [8,19,21,28], jackfruit [29], and other *Artocarpus* crop species [6,30,31]. Additionally, assembled and annotated a reference transcriptome of *A. altilis* and generated and analyzed 24 transcriptomes of breadfruit and its wild relatives to reveal signals of positive selection that may have resulted from local adaptation or natural selection [20]. Finally, a low coverage whole genome sequence has been published for *A. camansi* [32], but full genome sequences for jackfruit and breadfruit are still not available. Here, we report high quality annotated draft genome sequences for both jackfruit and breadfruit. Results help explain their energy-dense fruit composition and the underlying genes involved in sugar and starch metabolism.

## 2. Materials and methods

### 2.1. Sample collection, NGS Library construction, and sequencing

Genomic DNA was extracted from fresh leaves of *A. heterophyllus* (ICRAFF 11314) and *A. altilis* (ICRAFF 11315), grown at the World AgroForestry (ICRAF) campus in Kenya, using a modified CTAB method [33].

Extracted DNA was used to construct four paired-end libraries (170, 350, 500, and 800 bp) and four mate-pair libraries (2, 6, 10, and 20 Kb) following the standard protocols provided by Illumina (San Diego, USA). Subsequently, the sequencing was performed on a HiSeq 2000 platform (Illumina, San Diego, CA, USA) using a whole genome shotgun sequencing strategy. To improve the data quality, the poor quality reads were filtered using SOAPfilter (v2.2) [34]: (1) low-quality bases (Q = below 7 and 15 for *A. altilis* and *A. heterophyllus* respectively) were trimmed from both side of the reads; (2) removed reads with ≥30% low quality bases (quality score ≤ 15); (3) removed reads with ≥ 10% uncalled (“N”) bases; (4) removed reads with adapter contamination or PCR duplicates. (5) discarded reads with undersized insert sizes. Finally, more than 100x high-quality reads were obtained for each species (see Additional file: Table S1).

For transcriptome sequencing, the RNA was extracted from different tissues of *A. altilis* (various stages leaf, leaf bud, and roots) and *A. heterophyllus* (various stages of leaves, leaf bud, stem, bark, roots, germinated seed, and seedling). The RNA was extracted using the PureLink RNA Mini Kit (Thermo Fisher Scientific, Carlsbad, CA, USA) according to the manufacturer’s instructions. For each sample, RNA libraries were constructed by following the TruSeq RNA Sample Preparation Kit (Illumina, San Diego, CA, USA) manual, and were then sequenced on the Illumina HiSeq 2500 platform (paired-end, 100-bp reads), generating more than 47 Gb of sequence data for each species. Data were then filtered using a similar criterion as used to filter DNA NGS data, with a slight modification: (1) reads with ≥10% low-quality bases (quality score ≤15) were removed; and (2) reads with ≥5% uncalled (“N”) bases were removed (see Additional file: Table S2). All the transcriptome data from different tissues were compiled, and the combined version was used to check the completeness of the whole genome sequence assembly.

### 2.2. Evaluation of genome size

Clean reads of the paired-end libraries (170, 250 and 500 bp) were used to estimate the genome size by k-mer frequency distribution and heterozygosity analysis. The genome size was estimated based on the following formula:

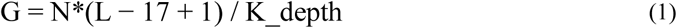

Where N represents the number of used reads, L represents the read length, K represents the k-mer value in the analysis and K_depth refers to the location of the main peak in the distribution curve [35]. The heterozygosity was evaluated by the GCE software [36].

### 2.3. De novo genome assembly

The *de novo* genome assembly tool, Platanus (Platanus, RRID:SCR_015531) [37], was used to construct the contigs and scaffolds in three steps: contig assembling, scaffolding and gap closing. In contig assembling, paired-end libraries ranging from 170 to 800 bp were used with the parameters “-d 0.5 -K 39 -u 0.1 -m 300”. In the scaffolding step, paired-end and mate-pair information were used to with parameters “-u 0.1”. Lastly, in the gap closing step, the paired-end reads were used with the parameters default. After the Platanus,we also using GapCloser for the gapclosing, GapCloser version 1.12 (GapCloser, RRID:SCR_015026) [34] with the parameters “-l 150 -t 32 -p 31” using pair-end libraries.

### 2.4. Genome assembly evaluation

The genome assembly completeness was assessed using BUSCO (Benchmarking Universal Single-Copy Orthologues), version 3.0.1 (BUSCO, RRID:SCR_015008) [38]. Next, the unigenes generated by Bridger software [39] from the transcriptome data of each species were aligned to the assembled genomes using BLAT (BLAT, RRID:SCR_011919) [40] with default parameters. Then, in order to confirm the accuracy of the assembly, some of the paired-end libraries (170, 250 and 350 bp) were aligned to the assembled genomes, and the sequencing coverage was calculated using SOAPaligner, version 2.21 (SOAPaligner/soap2, RRID:SCR_005503) [41].

We also calculated the GC content and average depth with 10 kb non-overlapping windows, the distribution of GC content indicated a relative pure single genome without contamination or GC bias (Additional file: Figure S3). Moreover, the GC content of *A. altilis* and *A. heterophyllus* genomes were also compared with three rosids species (*Fragaria vesca*, *Malus domestica*, and *Morus notabilis*).

### 2.5. Repeat annotation

Repetitive sequences were identified by using RepeatMasker (version 4-0-5) [42], with a combined library consisting of the Repbase library and a custom library obtained through careful self-training. The custom library was composed of three parts: the MITE, LTR and extensive library, which were constructed as described below.

First of all, the library of miniature inverted-repeat transposable elements (MITEs) was created by annotation using MITE-hunter [43] with default parameters. Secondly the library of long terminal repeat (LTR) was constructed using LTRharvest [44] integrated in Genometools (version 1.5.8) [45] with parameters “-minlenltr 100 -maxlenltr 6000 -mindistltr 1500 -maxdistltr 25000 -mintsd 5 -maxtsd 5 -similar 90 -vic 10” to detect LTR candidates in length of 1.5 kb to 25 kb, with two terminal repeats ranging from 100 bp to 6000 bp with >= 85% similarity. In order to improve the quality of the LTR library, we used several strategies to filter the candidate. As intact PPT (poly purine tract) or PBS (primer binding site) was necessary to define LTR, we subsequently use LTRdigest [46] with a eukaryotic tRNA library [47] to identify these features, and then removed the elements without appropriate PPT or PBS location. Subsequently, to remove contamination like local gene clusters and tandem local repeats, 50 bp of flanking sequences on both sides of each LTR candidates were aligned using MUSCLE (MUSCLE, RRID:SCR_011812) [48] with default parameters, if the identity >= 60%, the candidate was taken as a false positive and removed. LTR candidates which nested with other types of elements were also removed. Exemplars for the LTR library were extracted from the filtered candidates using a cutoff of 80% identity in 90% of sequence length. Furthermore, the regions annotated as LTRs and MITEs in the genome were masked, and then put into RepeatModeler version 1-0-8 RepeatModeler, RRID:SCR_015027) to predict other repetitive sequences for the extensive library.

Finally, the MITE, LTR and extensive libraries were integrated into the custom library, which was combined with Repbase library and then taken as the input for RepeatMasker to identify and classify repetitive elements genome-widely.

### 2.6. Gene prediction

Repetitive regions of the genome were masked before gene prediction. Based on the RNA, homologous and *de novo* prediction evidences the protein-coding genes were identified using the MAKER-P pipeline (version 2.31) [49]. For RNA evidence, the clean transcriptome reads were assembled into inchworms using Trinity version 2.0.6 [50], and then fed to MAKER-P as EST evidence. For homologous evidence, the protein sequences from four relative species in rosids (*F. vesca*, *M. domestica*, *M. notabilis*, *Prunus persica, Ziziphus jujuba*) were downloaded and provided as protein evidence.

For *de novo* prediction evidence, a series of training attempts were made to optimize different *ab initio* gene predictors. At first, a set of transcripts were generated by a genome-guided approach using Trinity with parameters “--full_cleanup --jaccard_clip --genome_guided_max_intron 10000 --min_contig_length 200”. The transcripts were then mapped back to the genome using PASA (version 2.0.2) [51] and a set of gene models with real gene characteristics (e.g. size and number of exons/introns per gene, features of spicing sites) was generated. The complete gene models were picked for training Augustus [52]. Genemark-ES (version 4.21) [53] was self-trained with default parameters. The first round of MAKER-P was run based on the evidence above with default parameters except “est2genome” and “protein2genome” set to “1”, yielding the only RNA- and protein-supported gene models. SNAP [54] was then trained with these gene models. Default parameters were used to run the second and final round of MAKER-P, producing final gene models.

Furthermore, non-coding RNA genes in the *A. altilis* and *A. heterophyllus* genomes were also annotated. BLAST tool was employed to search ribosomal RNA (rRNA) against *A.thaliana* rRNA database, or search microRNAs (miRNA) and small nuclear RNA (snRNA) against Rfam database (Rfam, RRID:SCR_004276)(release 12.0) [55]. tRNAscan-SE (tRNAscan-SE, RRID:SCR_010835) [56] was used to scan transfer RNA (tRNA) in the genome sequences.

### 2.7. Functional annotation of protein-coding genes

Functional annotation of protein-coding genes was based on sequence similarity and domain conservation by aligning translated coding sequences to public databases. The protein-coding genes were first queried against protein sequence databases, such as KEGG (KEGG, RRID:SCR_012773) [57], NR database (NCBI), COG [58], SwissProt and TrEMBL [59] for best-matches using BLASTP with an E-value cut-off of 1e-5. Secondly, InterProScan 55.0 (InterProScan, RRID:SCR_005829) [60] was used as an engine to identify the motif and domain-based on Pfam (Pfam, RRID:SCR_004726) [61], SMART (SMART, RRID:SCR_005026) [62], PANTHER (PANTHER, RRID:SCR_004869) [63], PRINTS (PRINTS, RRID:SCR_003412) [64] and ProDom (ProDom, RRID:SCR_006969) [65,66].

### 2.8. Ks-distribution analysis

The coding sequences and annotations for *Morus notabilis* were downloaded from the NCBI, reference RefSeq assembly accession GCF_000414095.1 [66]. The coding sequences and annotations for *Ziziphus jujube* [67] were downloaded from the Plaza4 database [68]. The headers of the. fasta files, as well as the 9^th^ columns of the. gff3 files were edited to make the datasets compatible with the software packages used for downstream analysis.

Ks-distribution analyses were performed, using the wgd-package [69]. For each species, the paranome was obtained by performing an all-against-all BlastP [70], with MCL clustering [71]. Codon multiple sequence alignment was done using MUSCLE [48]. Ks-distributions were constructed using codeml from the PAML4 package [72] and Fast-Tree [73] for inferring phylogenetic trees used in the node weighting procedure, other software used by the wgd. Thereafter, i-ADHoRe [74] was used to get anchor-point distributions and produce dot-plots. Lastly, Gausian mixture modes were fitted using 1 to 5 components.

### 2.9. One vs one synteny

One-vs-one synteny analysis was performed for pairs of the above-mentioned species, using the “work-flow 2” script that is part of the wgd-package [69].

### 2.10. Gene family construction

Protein and nucleotide sequences from *A. altilis*, *A. heterophyllus* and 7 species (*A. thaliana*, *F. vesca*, *M. domestica, M. notabilis, P. mume, P. persica, Z. jujuba*) were retrieved to construct gene families using OrthoMCL software [75] based on an all-versus-all BLASTP alignments with an E-value cutoff of 1e-5.

### 2.11. Phylogenetic analysis and divergence time estimation

We identified 486 single-copy genes in the 9 species, and subsequently used them to build the phylogenetic tree. Coding DNA sequence (CDS) alignments of each single-copy family were constructed following the protein sequence alignment with MUSCLE (MUSCLE, RRID:SCR_011812) [48]. The aligned CDS sequences of each species were then concatenated to a super gene sequence. The phylogenetic tree was constructed with PhyML-3.0 (PhyML, RRID:SCR_014629) [76] with the HKY85+gamma substitution model on extracted four-fold degenerate sites. Divergence time was calculated using the Bayesian relaxed molecular clock approach using MCMCTREE in PAML (PAML, RRID:SCR_014932) [72], based on the published calibration times (divergence between Arabidopsis thaliana and Rosales was 108-109 Mya, divergence between *P. mume* and *P. persica* was 24-72 Mya) [66]. The divergence time between *M. notabilis* and *Artocarpus* was predicted to be 61.8 (54.1-76.0) Mya (Figure 1A). Subsequently, to study gene gain and loss, CAFE (CAFE, RRID:SCR_005983) [77] was employed to estimate the universal gene birth and death rate λ (lambda) under a random birth and death model with the maximum likelihood method. The results for each branch of the phylogenetic tree were estimated (Figure 1A). Enrichment analysis on GO and pathway of genes in expanded families in the *Artocarpus* lineage were also calculated.

**Figure 1.**
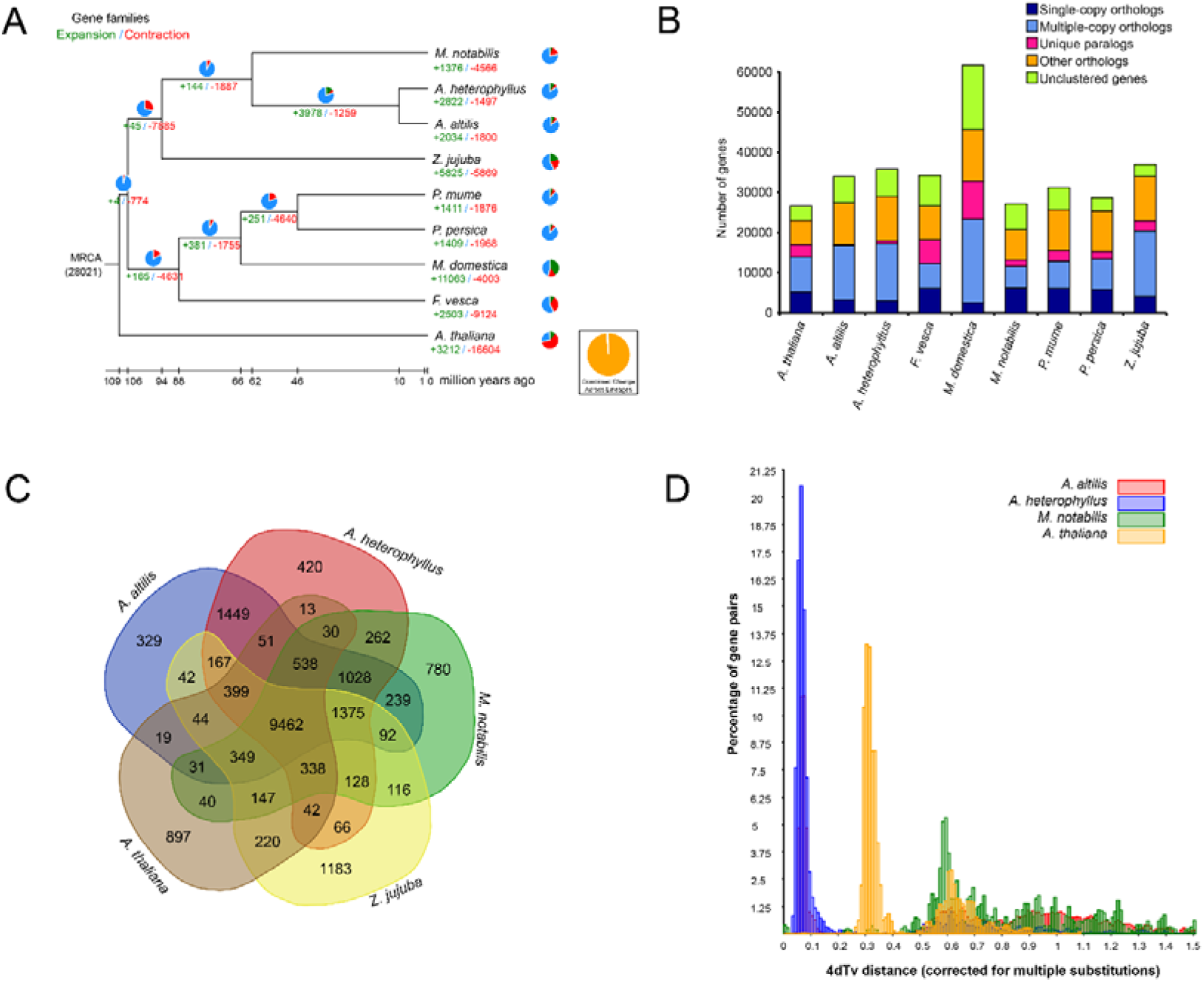
Phylogenetic and evolutionary analysis. (A, B) Gene conservation and gene family expansion and contraction in *A. heterophyllus* and *A. altilis*. The scale bar indicates 10 million years. The values at the branch points indicate the estimates of divergence time (mya), while the green numbers show the divergence time (million years ago, Mya), and the red nodes indicate the previously published calibration times. (C) The distribution of gene families among the model species and *Artocarpus* genus. (D) Distribution of 4DTv distance between collinearity gene pairs among *A. heterophyllus*, *A. altilis, M. notabilis* and *Arabidopsis thaliana*.

### 2.12. Identification of starch biosynthesis-related genes

Using the amino acid, starch biosynthesis-related genes in soybean as bait, we performed an ortholog search in *A. altilis, A. heterophyllus, M. notabilis, Z. jujuba, P. mume, P. persica, F. vesca, M. domestica and A. thaliana* (Figure 4).

## 3. Results and discussion

### 3.1. Genome sequencing and assembly

A total of eight libraries were constructed including four short-insert libraries (170 bp, 350 bp, 500 bp and 800 bp) and four mate-pair libraries (2 kb, 5 kb, 10 kb and 20 kb) for Illumina Hiseq2000 sequencing. In total, 273 Gb and 227 Gb of raw data was generated from *A. heterophyllus* and *A. altilis* respectively (Additional file: Table S1). We used the GCE software to evaluate the heterozygosity, and the results showed that the heterozygous ratio is 1.13% and 0.911% for *A. altilis* and *A. heterophyllus*, respectively. The K-mer distributions of *A. altilis* and *A. heterophyllus* showed two distinct peaks (Additional file: Figure S1, Figure S2), the first peaks was the heterozygous peak, the second peaks was the homozygous peak, where the second peak was confirmed as the main one for each of the species. Based on K-mer frequency methods [36], the *A. heterophyllus* and *A. altilis* genomes were estimated to be 1005 MB and 812 Mb, respectively (Additional file: Figure S1, Additional file: Table S3), the genome size of *A. altilis* and *A. heterophyllus* was relatively close to the genome size of species in the genus *Artocarpus* based on existing data in the C-values database, where 1C-value is 1.2 pg.

Using the SOAPdenovo2 program [41], all the *A. heterophyllus* high-quality reads were assembled into 108,267 scaffolds, totaling 982 Mb (Table 1). The N50s of contigs and scaffolds were 27 kb and 548 kb with longest being 255 kb and 3.1 Mb respectively (Table 1, Additional file: FigureS3). Similarly, for the *A. altilis*, the N50s of contigs and scaffolds were 17 kb and 1.5 Mb with longest being 174 kb and 7.4 Mb respectively (Table 1, Additional file: FigureS3). These results indicative of high quality of the assemblies for both the species. The GC content of the *A. heterophyllus* and *A. altilis* genomes were 32.9% and 32.3%, respectively. The GC depth graphs and distributions indicated there were no contamination in the genome assembly (Additional file: Figure S4).

**Table 1.**
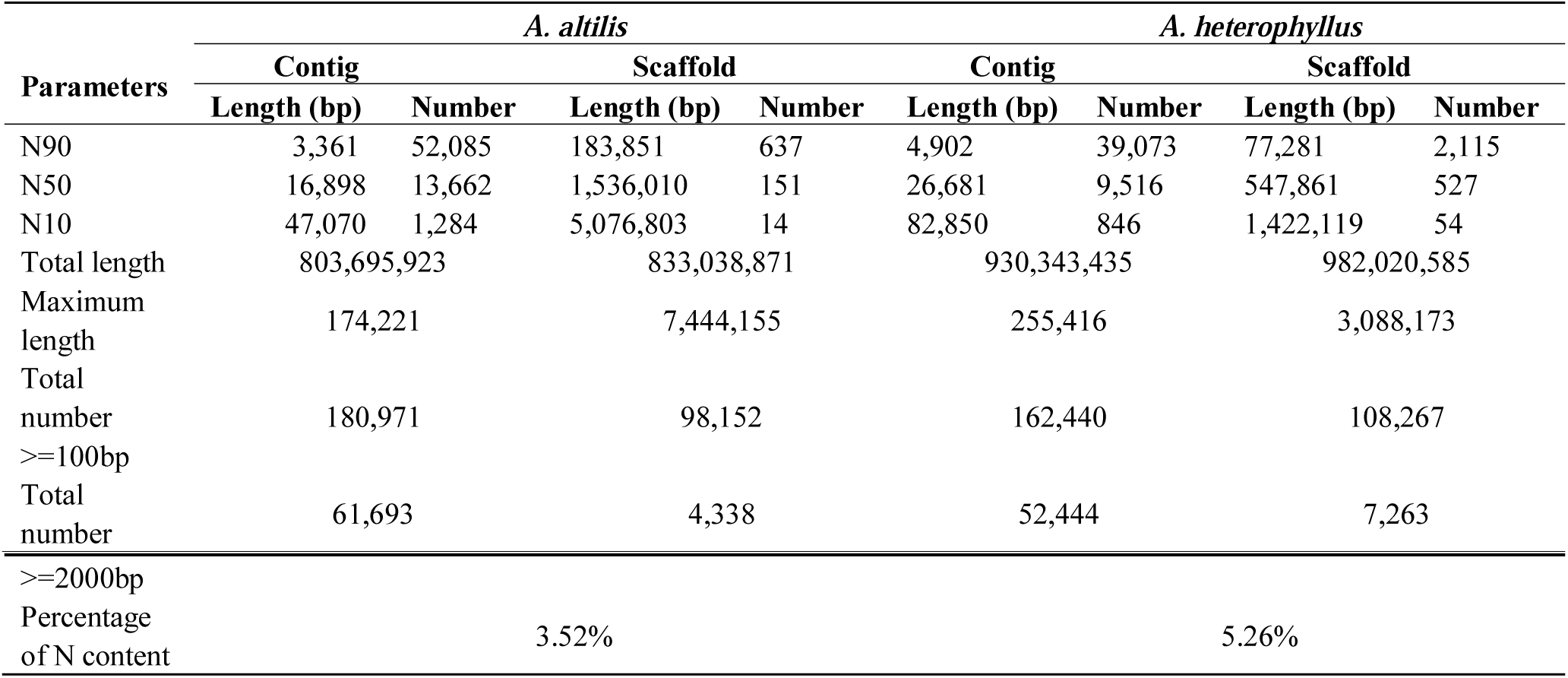
Statistics of the genome assembly of *A. altilis* and *A. heterophyllus*.

Evaluation of the quality and completeness of the draft genome assembly was done by the Benchmarking Universal Single-Copy Orthologs (BUSCO) data sets [38]. Of the total of 1440 BUSCO ortholog groups searched in the *A. heterophyllus* assembly, 932 (64.7%) BUSCO genes were “complete single-copy”, 437 (30.3%) were “complete duplicated”, 15 (1 %) were “fragmented”, and 56 (4 %) were “missing” (Tables 2). Similarly, in *A. altilis*, 988 (68.6%) BUSCO genes were “complete single-copy”, 383 (26.6%) were “complete duplicated”, 14 (~1 %) were “fragmented”, and 55 (3.8 %) were “missing” (Table 2), suggesting the high quality of the genome assembly. From the 1,440 core Embryophyta genes, 1,371 (95.20%) and 1,369 (95%) were identified in the *A. altilis* and *A. heterophyllus* assemblies, respectively (Table 2). We observed a significant difference in the number of duplicated core genes in *A. altilis* and *A. heterophyllus* [Table 2], which might be ascribed to the genome duplication in these species. The results also indicated that the assembly covered more than 90% of the expressed unigenes, suggesting the assembled genome covered a high percentage of expressed genes (Table 3). As expected, after the comparative GC content analysis the close peak positions showed *A. altilis*, *A. heterophyllus* and *M. notabilis* are closer than other species in GC content (Additional file: Figure S5).

**Table 2.** BUSCO evaluation of genome assembly of *A. altilis* and *A. heterophyllus*.

| BUSCOs | *A. altilis* |  | *A. heterophyllum* |  |
| --- | --- | --- | --- | --- |
|  | N | P (%) | N | P (%) |
| Complete BUSCOs | 1371 | 95.20 | 1369 | 95.00 |
| Complete single-copy | 988 | 68.60 | 932 | 64.70 |
| Complete duplicated | 383 | 26.60 | 437 | 30.30 |
| Fragmented | 14 | 1.00 | 15 | 1.00 |
| Missing | 55 | 3.80 | 56 | 4.00 |

Abbreviation: BUSCO, Benchmarking universal single-copy orthologs; N, number; P, percentage of complete BUSCOs compared to the total BUSCOs

**Table 3.**
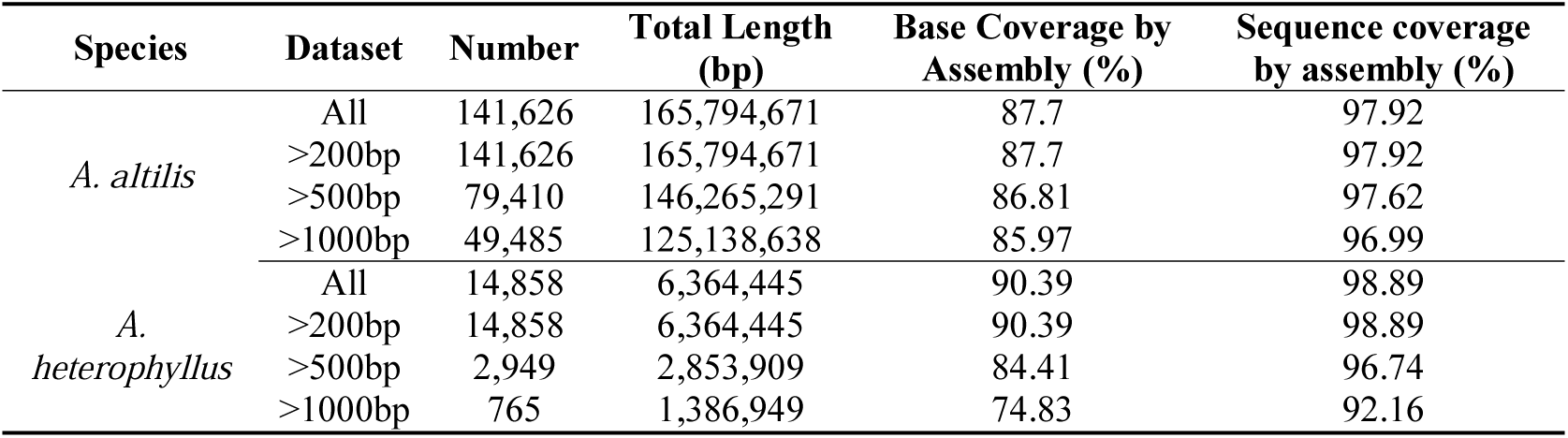
The gene coverage based on transcriptome data.

### 3.2. Gene annotation

A combination of *de novo* and homology-based methods (using transcript data as evidence) were used to identify repeat sequences. We found that up to 51.01% of the *A. heterophyllus* and 52.04% of the *A. altilis* assembled sequences were repeat sequences, comprised mostly of transposable elements and tandem repeats. Interestingly the amounts of these elements were higher than what is observed in orange (20%, 367 Mb) [78], peach (29.6%, 265 Mb) [79], pineapple (38.3%, 526 Mb) [80] and others (Table 4). This is consistent with the finding that bigger fruit tree genomes often retained higher percentages of repetitive elements compared to the smaller fruit tree genomes [81]. Among the repetitive sequences, 36.99% and 45.95% were of the long terminal repeat (LTR) type, respectively (Table 4), indicating LTRs are the most abundant transposable elements in *A. heterophyllus* and *A. altilis* genomes.

**Table 4.** Classification of predicted transposable elements in the genome of *A. altilis* and *A. heterophyllus*.

| Repeat type | *A. altilis* |  | *A. heterophyllum* |  |
| --- | --- | --- | --- | --- |
|  | % in genome | Length (bp) | % in genome | Length (bp) |
| SINE | 0 | 1,187 | 0.03 | 384,983 |
| LINE | 0.14 | 1,214,650 | 0.99 | 9,775,316 |
| LTR | 45.95 | 382,841,531 | 36.99 | 363,293,617 |
| DNA | 2.95 | 24,608,939 | 3.76 | 36,982,825 |
| Satellite | 0 | 34,585 | 0.3 | 3,001,478 |
| Simple repeat | 0.03 | 253,818 | 0.04 | 485,582 |
| Unknow | 5.4 | 45,013,282 | 12.23 | 120,128,962 |
| Total | 52.04 | 433,486,547 | 51.01 | 500,968,186 |

Using a comprehensive annotation strategy, we annotated a total of 35,858 *A. heterophyllus* genes and 34,010 *A. altilis* genes (Table 5). This was close to the number of genes (39,282) predicted in *Dimocarpus longan*, an exotic round to oval Asian fruit [81]. The average *A. heterophyllus* gene length was 3472.22 bp, the average length of the coding sequence (CDS) was 1241.48 bp, and the average number of exons per gene was 5.48 (Table 5, Additional file: Figure S6). We predicted a total of 466 rRNA, 159 miRNA, 1,554 snRNA genes and 713 tRNA in *A. altilis*, and a total of 2,706 rRNA, 168 miRNA, 1,005 snRNA genes and 689 tRNA in *A. heterophyllus* (Table 6). Of 35,858 *A. heterophyllus* protein-coding genes, 35,076 (97.82%) had Nr homologs, 34,968 (97.52%) had TrEMBL homologs, 27,632 (77.06%) had InterPro homologs and 27,741 (77.36%) had SwissProt homologs (Table 7). Similar to *A. heterophyllus*, the average *A. altilis* gene size was 3545.36 bp, the average length of the CDS was 1252.56 bp, and the average number of exons per gene was 5.50 (Table 5). Of 34,010 *A. altilis* protein-coding genes, 33,353 (98.07%) had Nr homologs, 33,240 (97.74%) had TrEMBL homologs, 26,422 (77.69%) had InterPro homologs and 26,689 (78.47%) had SwissProt homologs (Table 7). BUSCO evaluation showed that more than 89% of 1440 core genes were complete, suggesting an acceptable gene annotation for *A. altilis* and *A. heterophyllus* genomes (Additional file: Table S4).

**Table 5.** Statistics of gene models of *A. altilis*, *A. heterophyllus* and other species in Rosids.

|  | *A. altilis* | *A. heterophyllum* | *F. vesca* | *M. domestica* | *M. notabilis* | *P. persica* | *Z. jujuba* |
| --- | --- | --- | --- | --- | --- | --- | --- |
| Protein-coding gene number | 34,010 | 35,858 | 34,301 | 61,721 | 27,085 | 28,701 | 37,526 |
| Mean gene length (bp) | 3,545.36 | 3,472.22 | 2,824.55 | 2,692.45 | 2,866.82 | 2,464.79 | 3,313.54 |
| Mean cds length (bp) | 1,252.56 | 1241.48 | 1174.73 | 1141.42 | 1086.85 | 1210.76 | 1352.96 |
| Mean exons per gene | 5.49 | 5.48 | 5.05 | 4.82 | 4.6 | 4.97 | 5.5 |
| Mean exon length (bp) | 227.75 | 226.46 | 232.51 | 236.7 | 236.35 | 243.59 | 245.98 |
| Mean intron length (bp) | 509.56 | 497.69 | 407.11 | 405.79 | 494.64 | 315.84 | 435.66 |

**Table 6.**
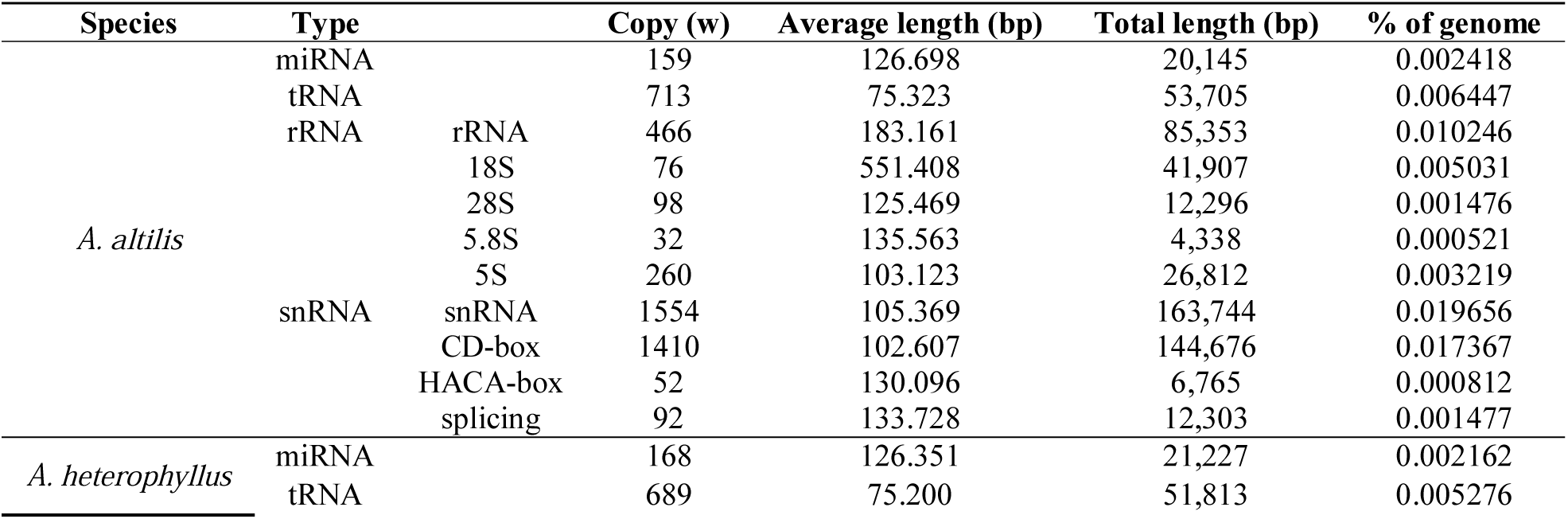
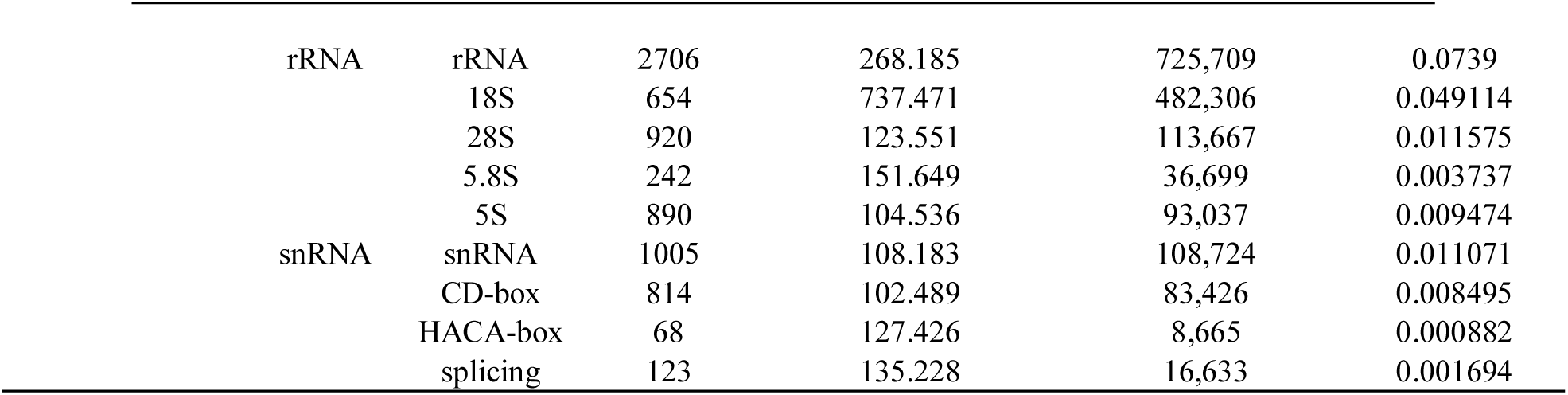
Annotation of non-coding RNA genes in the *A. altilis* and *A. heterophyllus* genomes.

**Table 7.** Statistics of functional annotation of protein-coding genes in the *A. altilis* and *A. heterophyllus genomes*.

| Values | *A. altilis* |  | *A. heterophyllus* |  |
| --- | --- | --- | --- | --- |
|  | Number | Percentage | Number | Percentage |
| Total | 34,010 | 100% | 35,858 | 100% |
| Nr | 33,353 | 98.07% | 35,076 | 97.82% |
| Swissprot | 26,689 | 78.47% | 27,741 | 77.36% |
| KEGG | 24,860 | 73.10% | 25,804 | 71.96% |
| COG | 12,875 | 37.86% | 13,408 | 37.39% |
| TrEMBL | 33,240 | 97.74% | 34,968 | 97.52% |
| Interpro | 26,422 | 77.69% | 27,632 | 77.06% |
| GO | 17,428 | 51.24% | 18,336 | 51.14% |
| Overall | 33,394 | 98.19% | 35,109 | 97.91% |
| Unannotated | 616 | 1.81% | 749 | 2.09% |

### 3.3. Gene family evolution and comparison

Orthologous clustering analysis was conducted with the *A. altilis* and *A. heterophyllus* genomes following comparison with seven other plant genomes: *A. thaliana*, *F. vesca*, *M. domestica, M. notabilis, P. mume, P. persica, Z. jujuba*. A Venn diagram shows that *A. altilis*, *A. heterophyllus*, *A. thaliana*, *M. notabilis*, *Z. jujuba* contain a core set of 9462 gene families in common, there were 1028 orthologous families shared by three Moraceae species, while 329 gene families containing 515 genes were specific to *A. altilis*, and 420 gene families containing 907 genes were specific to *A. heterophyllus*. (Figure 1C).

Of the 35,845 protein-coding genes in the *A. heterophyllus* genome, 28,969 were grouped into 15,768 gene families (of which 242 were *A. heterophyllus*-unique families) (Figure 1B, Additional file: Table S5). Of the 33,986 *A. altilis* protein-coding genes, 27,354 were grouped into 15,614 gene families (of which 136 were *A. altilis*-unique families) (Figure 1B, Additional file: Table S5).

Phylogenetic analysis showed that *A. heterophyllus* and *A. altilis* were more closely related to mulberry than to Jujube (Figure 1A), further supporting a previous phylogeny of *Artocarpus* [2]. CAFE [77] was used to identify gene families that had potentially undergone expansion or contraction. We found a total of 2,822 expanded gene families and 1,497 contracted families in *A. heterophyllus*, as well as 2034 expanded and 1800 contracted families in *A. altilis* (Figure 1A). The genes in the expanded and contracted families were assigned to Kyoto Encyclopedia of Genes and Genomes (KEGG) pathways [82]. The *A. heterophyllus*-expanded gene families were remarkably enriched in metabolism related pathways/functions, including starch and sucrose metabolism (ko00500, P=0.003), glycan degradation (ko00511, P=0.007), glycolysis/gluconeogenesis (ko00010, P=0.016) and others (Additional file: Table S7). KEGG enrichment analysis of *A. altilis* revealed that pathways associated with photosynthesis, such as Carbon fixation in photosynthetic organisms (ko00710, P=0.017), Other types of O-glycan biosynthesis (ko00514, P=0.018) and Photosynthesis (ko00195, P=0.006) were particularly enriched (Additional file: Table S7).

In order to determine whether there is any evidence for whole genome duplications in *A. heterophyllus* and *A. altilis*, the distance–transversion rates at 4-fold degenerate sites (4DTv) was calculated (Figure 1D, Additional file: Figure S7). Two 4DTv values that peaked at 0.07 and 0.08 for orthologs between *A. heterophyllus*, and between *A. altilis* respectively, which highlighted the recent whole-genome duplication of these two species. The results of the Ks distributions mostly corroborate the findings of the 4DTv analysis. The results suggest that the whole genome duplication event was shared by *Arthrocarpus altilis* and *Arthrocarpus heterophylus*. Their divergence is recent, as suggested by the overlap of their WGD peaks (Figure 2), meaning that they have equal substitution, duplication and loss rates. Thus, for further analysis (one-vs-one synteny with the close relatives *M. notabilis* and *Z. jujube)*, only *A. altilis* was used. These results suggest that the *Athrocarpus* genome duplication event occurred after divergence from the common ancestor they share with *M. notabilis* (Figure 3), thus between 62 and 10 MYA.

**Figure 2.**
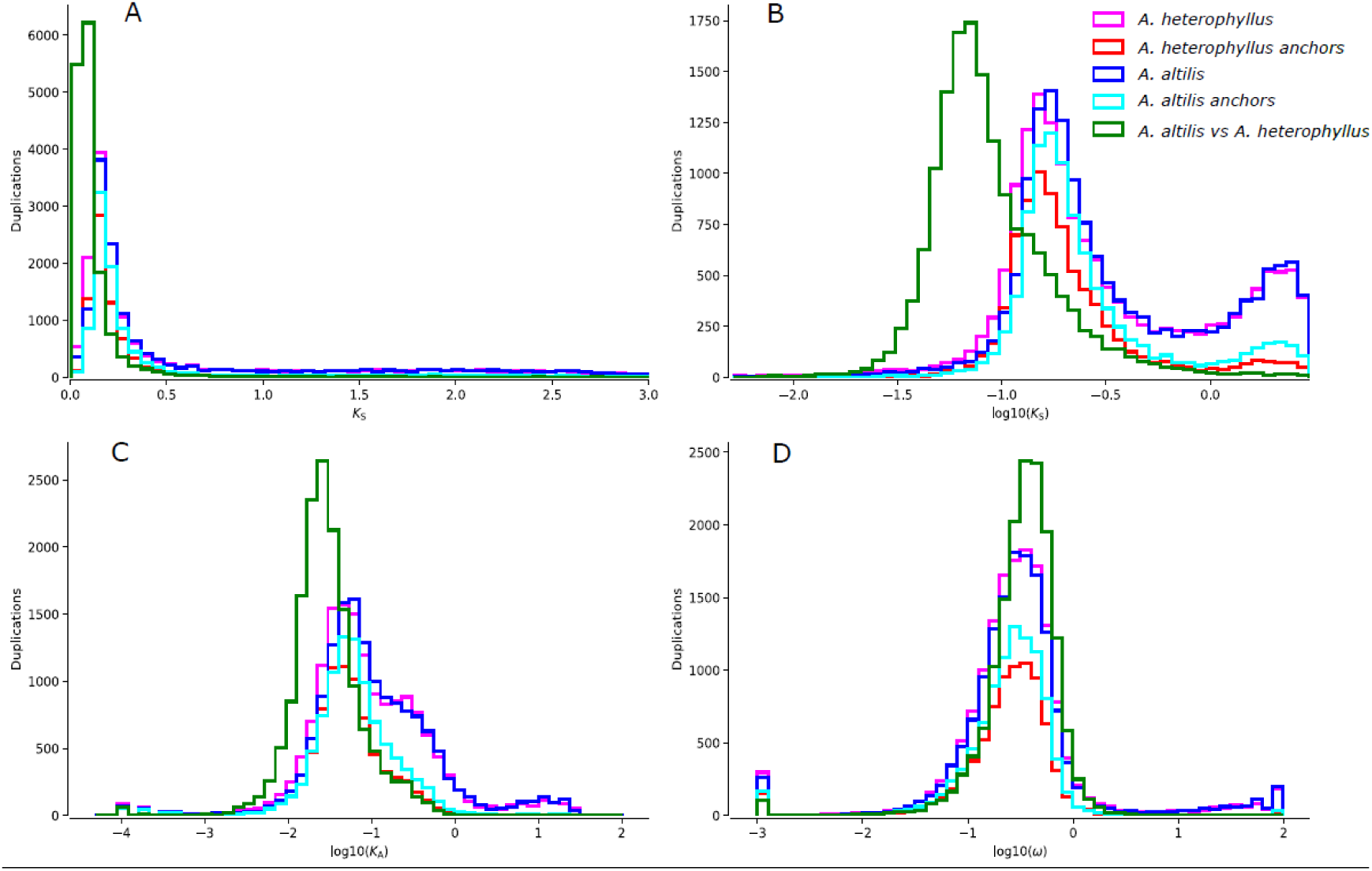
A graph showing the *A. heterophyllus* Ks distributions (light pink) and the Ks distributions of its anchor pairs (red), *A altilis* Ks distributions (blue) and the distributions of its anchor pairs (light blue) overlaid with the Ks distributions of the one-to-one orthologs of *A. heterophyllus* and *A. altilis* (green). Figure 2B, log transformed version of 2A. Figures 2C and 2D are the log transformed Ka and ω distributions respectively.

**Figure 3.**
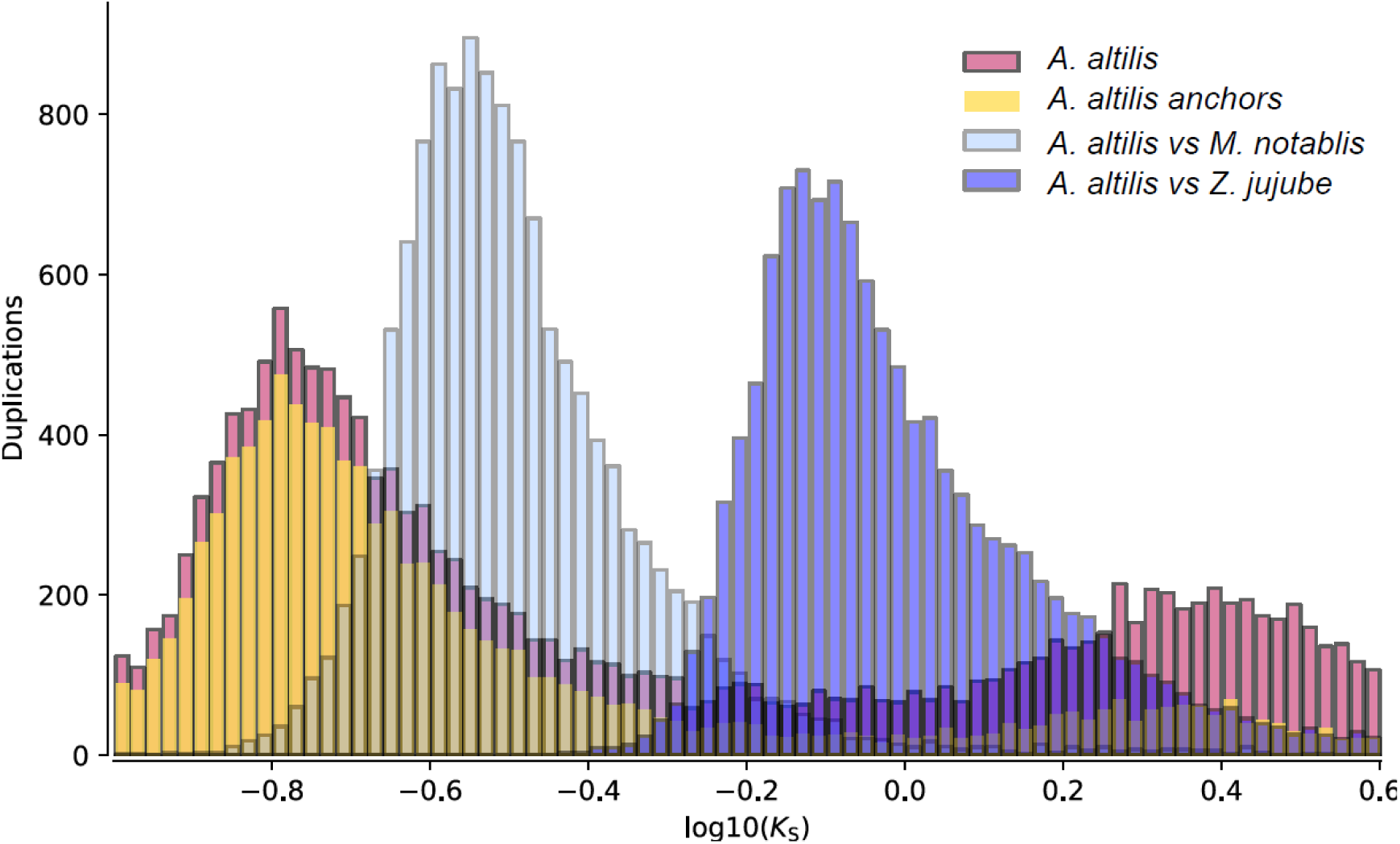
Ks distribution (dark pink) and anchor pair Ks distribution (yellow) of *A. altilis* in overlay with the results of whole paranome distributions between *A. altilis* and *M. notabilis* (light blue) and *A. altilis* and *Z. jujube* (dark blue).

### 3.4. Gene family expansion and tissue specific expression of starch synthesis related genes

The copy number of starch synthesis related genes were compared between *A. heterophyllus*, *A. altilis*, closely related species, as well as some other starch-rich plant species (Figure 4). We observed a remarkable copy number expansion of the *UGD1* gene in *A. heterophyllus* compared with the other species. The enzyme encoded by *UGD1*, catalyzes the conversion of Glucose-1-P into UDP-GlcA, thereby stalling the starch synthesis process [83] (Figure 4). Interestingly, the tissue-specific expression pattern of *UGD1* contrasts with other starch synthesis genes in *A. heterophyllus* (Figure 5A). For instance, in *A. heterophyllus* there is a suppression of UDPG transcription in the stem, while the other starch biosynthesis genes are activated. However, differential expression of *UGD1* was not shown in *A. altilis* (Figure 5B). This unusual expression pattern of *UGD1* as well as the gene copy number expansion might lead to the failure of starch accumulation in *A. heterophyllus* rather than *A. altilis*. But this needs to be further validated by real time-qPCR for confirmation of the tissue specific expression. For the GO enrichment, expansion of gene families were related to small molecule binding or single organism signaling (Additional file: Table S6) in *A. altilis*. Moreover, there were some expansion of gene families related to molecule binding, reproductive process and cellular response to stimulus in *A. heterophyllus*. Gene families belonging to expanded pathways in *A. altilis* were mainly related to plant-pathogen interaction, Lysine biosynthesis or photosynthesis. In contrast the gene families that were expanded in *A. heterophyllus* belonged to pathways involving secondary metabolite biosynthesis, phenylpropanoid biosynthesis and fatty acid metabolism. In contrast, the Biosynthesis of secondary metabolites, Phenylpropanoid biosynthesis and Fatty acid metabolism were enriched in the expanded gene families in *A. heterophyllus*. (Additional file: Table S7).

**Figure 4.**
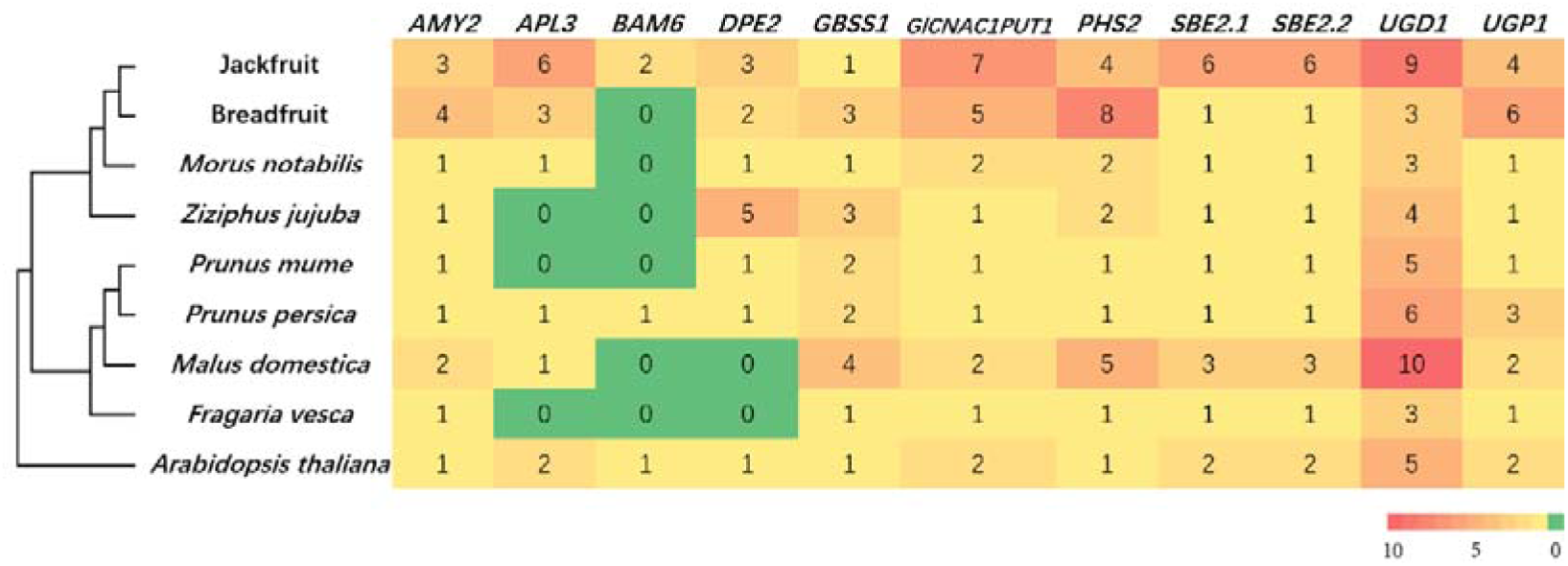
Copy number expansion of starch synthesis related genes in *A. heterophyllus* and *A. altilis*.

**Figure 5.**
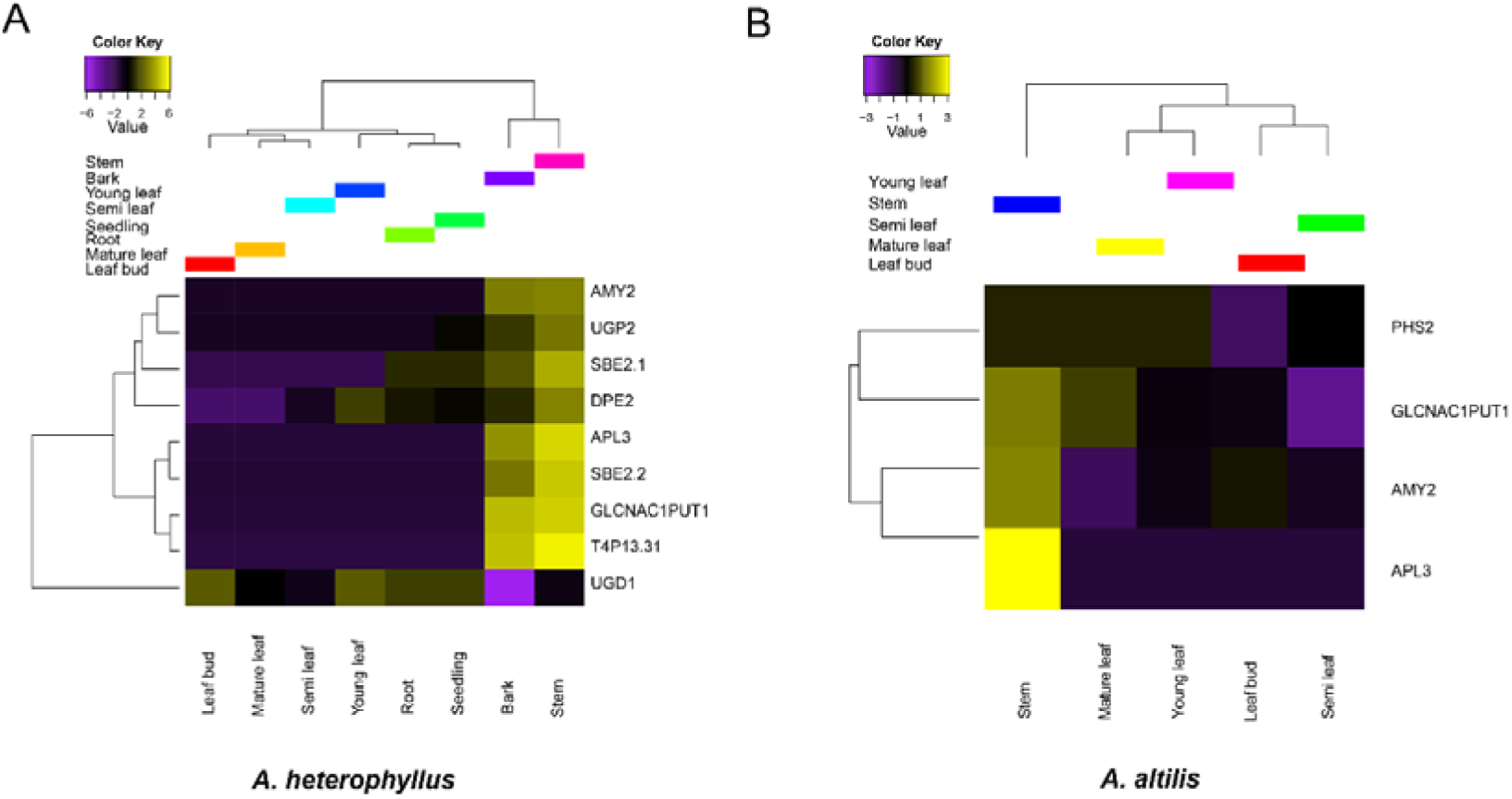
Tissue specific expression of starch synthesis related genes in *A. heterophyllus* and *A. altilis*.

## 4. Conclusion

Here, we report the genomes of jackfruit (*A. heterophyllus*) and breadfruit (*A. altilis*). The publication of these high-quality draft genomes and annotations may provide plant breeders and other researchers with useful information regarding trait biology and their subsequent improvement. In particular, we highlight genes unique to *A. heterophyllus* and *A. altilis* due to their high sugar and starch content (respectively), which are desirable characteristics in these edible plants. The information provided in the draft genome annotations can be used to accelerate genetic improvement of these crops. The availability of these genomes on the AOCC ORCAE platform (https://bioinformatics.psb.ugent.be/orcae/aocc) will enable various stakeholders to access and improve the annotations of these genomes.

## Supplementary Materials

Figure S1: K-mer (K=17) analysis of the two genomes, Figure S2: Distribution of sequencing depth of the assembly data, Figure S3: Distribution of the length and number of the scaffold in two species, Figure S4: The GC content, Figure S5: Comparison of GC content across closely related species, Figure S6: Statistics of gene models in *A. altilis, A. heterophyllus, F. vesca, M. domestica, M. notabilis, Prunus. persica* and *Ziziphus. Jujube*, Figure S7: The collinearity between two species, Table S1. Statistics of the raw and clean data of DNA sequencing, Table S2. Summary statistics of the transcriptome data, Table S3. Estimation of the genome size based on K-mer statistics, Table S4. BUSCO evaluation of the annotated protein-coding genes in *A. altilis* and *A. heterophyllus*, Table S5. Analysis of gene families of different species, Table S6. Enriched GO terms (level 3) of genes in families with expansion, Table S7. Enriched pathways of genes in families with expansion.

## Author contributions

Conceptualization, Robert Kariba, Xun Xu, Allen Van Deynze, Xin Liu and Huan Liu; Data curation, Sunil Kumar Sahu, Min Liu, Bo Song, Shu-Min Kao, Nyree J.C. Zerega and Yves Van de Peer; Formal analysis, Sunil Kumar Sahu, Min Liu, Anna Yssel and Sanjie Jiang; Funding acquisition, Prasad S. Hendre, Xun Xu, Huanming Yang, Xin Liu and Huan Liu; Investigation, Sunil Kumar Sahu, Samuel Muthemba and Prasad S. Hendre; Methodology, Min Liu; Project administration, Xun Xu, Huanming Yang, Xin Liu and Huan Liu; Resources, Robert Kariba, Samuel Muthemba, Ramni Jamnadass, Shu-Min Kao, Nyree J.C. Zerega, Yves Van de Peer, Jonathan Featherston and Huan Liu; Software, Min Liu; Supervision, Prasad S. Hendre, Huanming Yang, Allen Van Deynze, Yves Van de Peer, Xin Liu and Huan Liu; Validation, Min Liu; Visualization, Anna Yssel; Writing – original draft, Sunil Kumar Sahu; Writing – review & editing, Min Liu, Anna Yssel, Robert Kariba, Sanjie Jiang, Bo Song, Samuel Muthemba, Prasad S. Hendre, Ramni Jamnadass, Shu-Min Kao, Nyree J.C. Zerega, Xun Xu, Huanming Yang, Allen Van Deynze, Yves Van de Peer, Jonathan Featherston, Xin Liu and Huan Liu.

## Availability of supporting data

The genome and transcriptome data are deposited in the CNGB Nucleotide Sequence Archive (CNSA: http://db.cngb.org/cnsa; accession number CNP0000715, CNP0000486), and all the annotations are also available via AOCC ORCAE platform (https://bioinformatics.psb.ugent.be/orcae/aocc)

