## Additional file for "Draft Genomes of two *Artocarpus* plants, Jackfruit (*A. heterophyllus*) and Breadfruit (*A. altilis*)"

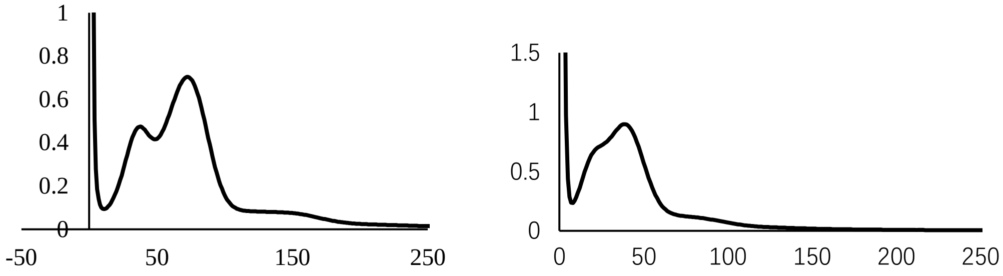


**A** **B**

**Figure S1: K-mer (K=17) analysis of the two genomes.** The X-axis is depth; the y-axis represents the frequency, A: *A. altilis*, B: *A. heterophyllus*. In the A, the left peak was the heterozygous peak, the right peak was the homozygous peak. Same as B.

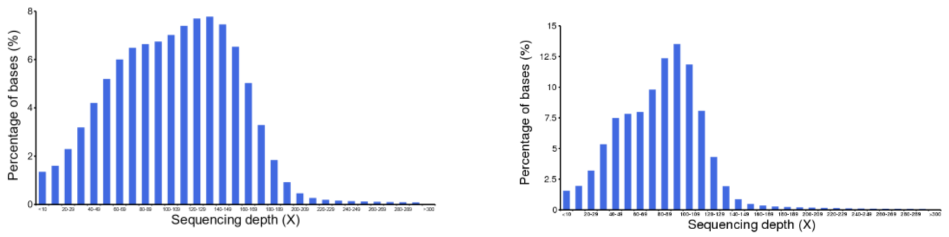


A B

**Figure S2: Distribution of sequencing depth of the assembly data.** The X-axis is the depth and the y-axis show the percentage of bases at each depth. The results show that <1% of bases had a sequencing depth less than 10. and two peaks demonstrate the genome heterozygosity. A: *A. altilis*, B: *A. heterophyllus*.

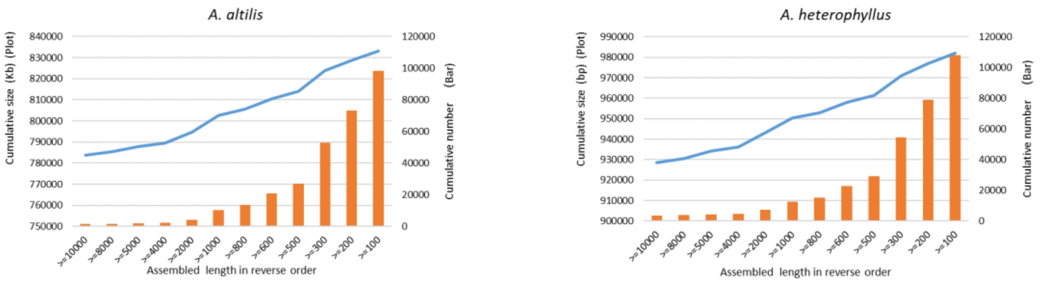


Figure S3: Distribution of the length and number of the scaffold in two species. The blue lines showes the size, and the orange bar shows the number.

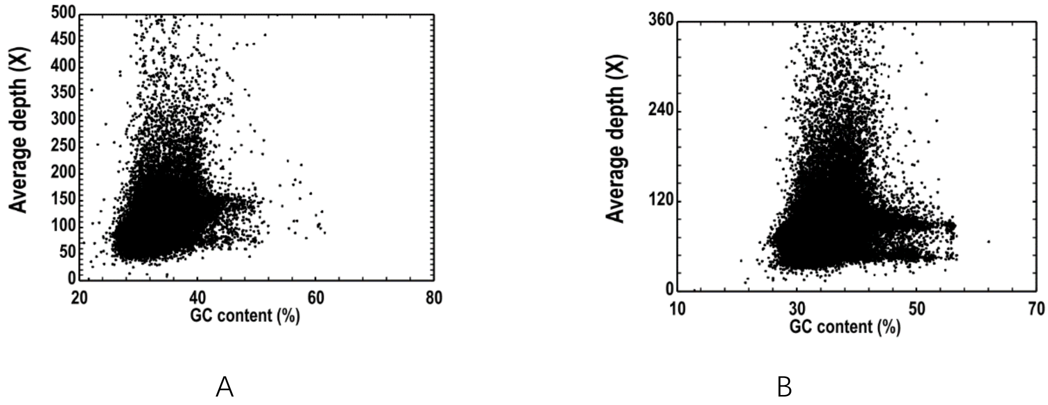


**Figure S4: The GC content.** The GC content and the average depth were calculated from 10 kb non-overlapping sliding windows. The distribution pattern of GC content indicates a relative pure single genomic sample without contamination and no GC bias, but repeat. A: *A. altilis*, B: *A. heterophyllus*.

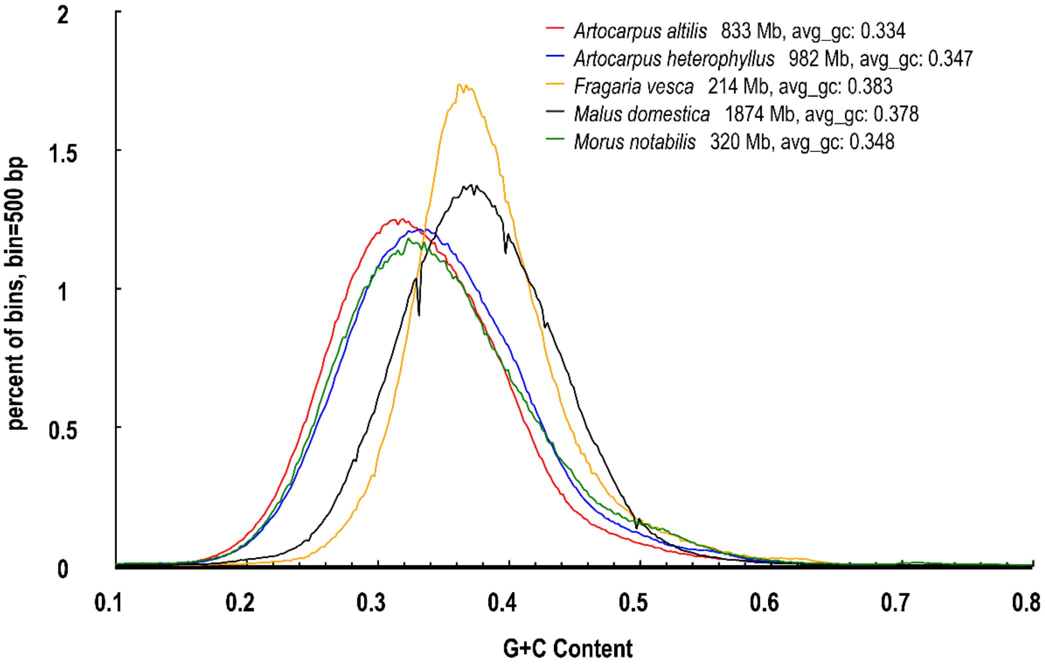


**Figure S5: Comparison of GC content across closely related species.** The *A. altilis*, *A. heterophyllus* and *M. notabilis* belong to the same family, so the show the same peaks of GC content.

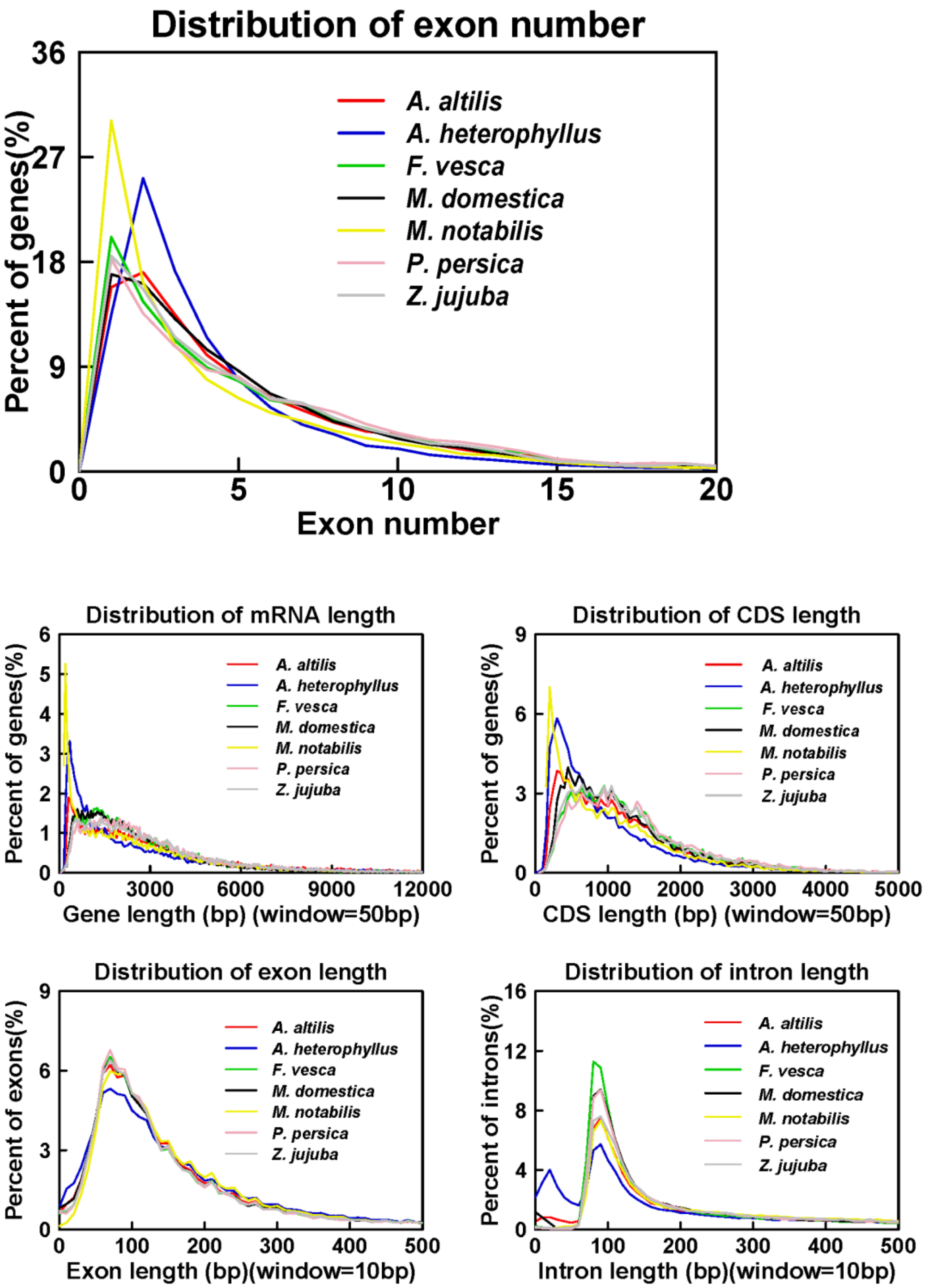


**Figure S6: Statistics of gene models in *A. altilis, A. heterophyllus, F. vesca, M. domestica, M. notabilis, Prunus persica and Ziziphus jujuba.***

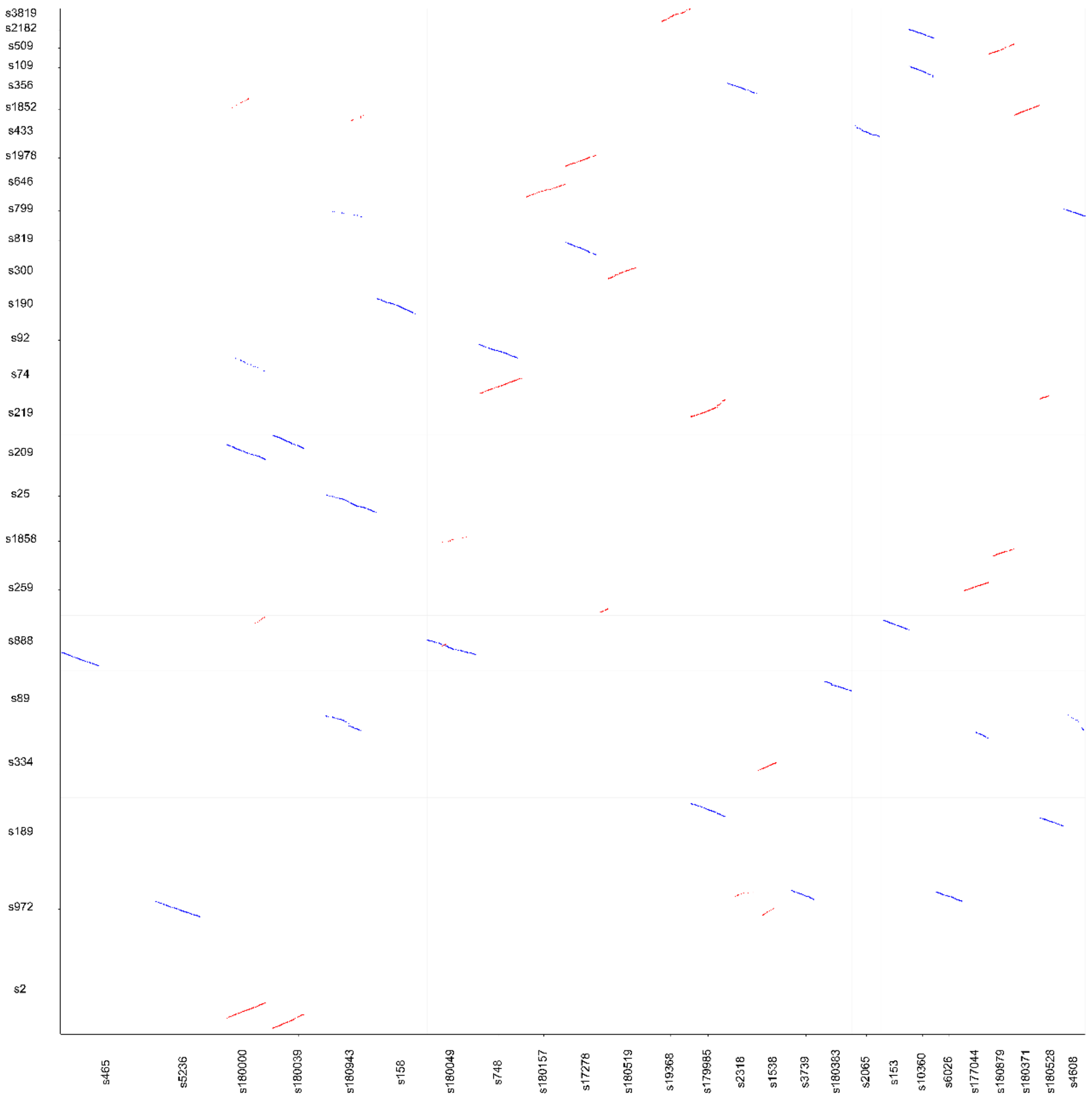


**Figure S7: The collinearity between two species.** The X-axis is the *A. heterophyllus*; Y-axis is the *A. altilis*.

**Table S1.** Statistics of the raw and clean data of DNA sequencing. Clean data were obtained by filtering raw data as described in the article. The sequencing depth calculated based on a genome size of 811, 1005 Mb of *A. altilis* and *A. heterophyllus*, respectively.

| Species | Library insert size (bp) | Read length (bp) | Raw data | | | Clean data | | |
|---|---|---|---|---|---|---|---|---|
| | | | base number (bp) | Reads number (bp) | Depth (X) | base number (bp) | Reads number (bp) | Depth (X) |
| *A. altilis* | 170 | PE100 | 27,997,546,200 | 279,975,462 | 34.51 | 24,457,319,840 | 257,445,472 | 30.15 |
| | 350 | PE100 | 37,756,156,600 | 377,561,566 | 46.54 | 32,314,250,380 | 323,142,504 | 39.83 |

| | | | | | | | | |
|---|---|---|---|---|---|---|---|---|
| | 500 | PE100 | 17,016,336,900 | 170,163,369 | 20.98 | 14,394,991,000 | 143,949,910 | 17.74 |
| | 800 | PE100 | 27,820,793,600 | 278,207,936 | 34.29 | 23,427,277,780 | 234,272,778 | 28.88 |
| | 2000 | PE100 | 29,259,701,000 | 292,597,010 | 36.07 | 11,676,799,030 | 116,767,990 | 14.39 |
| | 6000 | PE100 | 26,886,185,600 | 268,861,856 | 33.14 | 10,018,166,480 | 100,181,665 | 12.35 |
| | 10000 | PE100 | 34,525,749,400 | 345,257,494 | 42.56 | 10,295,921,210 | 102,959,212 | 12.69 |
| | 20000 | PE100 | 25,992,313,200 | 259,923,132 | 32.04 | 3,530,699,890 | 35,306,999 | 4.35 |
| | Total | | 227,254,782,500 | 2,272,547,825 | 280.13 | 130,115,425,610 | 1,314,026,530 | 160.39 |
| *A. heterophyllus* | 170 | PE100 | 38,595,699,600 | 385,956,996 | 38.38 | 33,017,200,030 | 330,172,000 | 32.83 |
| | 350 | PE100 | 32,428,208,400 | 324,282,084 | 32.25 | 27,992,453,380 | 279,924,534 | 27.84 |
| | 500 | PE100 | 14,831,029,800 | 148,310,298 | 14.75 | 12,425,626,520 | 124,256,265 | 12.36 |
| | 800 | PE100 | 22,618,262,000 | 226,182,620 | 22.49 | 19,410,382,710 | 194,103,827 | 19.30 |
| | 2000 | PE100 | 76,821,363,600 | 768,213,636 | 76.39 | 36,971,268,180 | 369,712,682 | 36.77 |
| | 6000 | PE100 | 32,718,039,600 | 327,180,396 | 32.54 | 14,483,081,550 | 144,830,816 | 14.40 |
| | 10000 | PE100 | 22,875,623,600 | 228,756,236 | 22.75 | 4,204,951,180 | 42,049,512 | 4.18 |
| | 20000 | PE100 | 32,777,398,400 | 327,773,984 | 32.59 | 4,311,498,050 | 43,114,981 | 4.29 |
| | Total | | 273,665,625,000 | 2,736,656,250 | 272.14 | 152,816,461,600 | 1,528,164,616 | 151.96 |

**Table S2.** Summary statistics of the transcriptome data.

| | | Raw data | | Clean data | | |
|---|---|---|---|---|---|---|
| Species | Abbreviation | Base number (bp) | Reads number (bp) | Base number (bp) | Reads number (bp) | Sample |
| *A. altilis* | AALBd | 4,629,731,282 | 19,131,121 | 611,897,508 | 2,886,309 | Leaf bud |
| | AAYL | 11,655,613,464 | 48,163,692 | 1,117,701,948 | 5,272,179 | Young leaf |

| | | | | | | |
|---|---|---|---|---|---|---|
| | AASL | 10,178,838,956 | 42,061,318 | 1,024,350,292 | 4,831,841 | Semi mature leaf |
| | AAML | 19,051,406,682 | 78,724,821 | 2,635,339,776 | 12,430,848 | Mature leaf |
| | AAST | 8,529,531,032 | 35,245,996 | 916,693,088 | 4,324,024 | stem |
| | Total | 54,045,121,416 | 223,326,948 | 6,305,982,612 | 29,745,201 | |
| *A. heterophyllus* | AHLB | 1,174,736,970 | 9,708,570 | 20,728,520 | 180,248 | Leaf bud |
| | AHYL | 11,510,346,914 | 95,126,834 | 194,000,860 | 1,686,964 | Young leaf |
| | AHML | 928,769,616 | 7,675,782 | 66,734,270 | 580,298 | Mature leaf |
| | AHSL | 13,289,054,416 | 109,826,896 | 200,927,310 | 1,747,194 | Semi mature leaf |
| | AHRT | 2,612,876,178 | 21,594,018 | 232,498,720 | 2,021,728 | Roots |
| | AHSDL | 12,200,805,826 | 100,833,106 | 1,097,277,560 | 9,541,544 | Seedling |
| | AHS | 6,078,062,078 | 50,231,918 | 1,001,884,60 | 871,204 | Stem |
| | Total | 47,794,651,998 | 394,997,124 | 1,812,167,240 | 16,629,180 | |

**Table S3.** Estimation of the genome size based on K-mer statistics.

| Species | Kmer value | Kmer number | Peak depth(X) | Genome size (Mb) | Used bases (Gb) | Used reads (Mb) | Depth (X) |
|---|---|---|---|---|---|---|---|
| *A. altilis* | 17 | 59,221,955,980 | 73 | 811.26 | 70.08 | 678.89 | 44.96 |
| *A. heterophyllus* | 17 | 39,218,751,230 | 39 | 1005.61 | 46.29 | 442.23 | 46.03 |

**Table S4.** BUSCO evaluation of the annotated protein-coding genes in *A. altilis* and *A. heterophyllus.*

| BUSCOs | *A. altilis* | | *A. heterophyllus* | |
|---|---|---|---|---|
| | N | P(%) | N | P(%) |
| Complete BUSCOs | 1,319 | 91.6 | 1,288 | 89.5 |
| Complete single-copy | 977 | 67.8 | 885 | 61.5 |
| Complete duplicated | 342 | 23.8 | 403 | 28 |
| Fragmented | 32 | 2.2 | 30 | 2.1 |

| | | | | |
|---|---|---|---|---|
| Missing | 89 | 6.2 | 122 | 8.4 |

**Table S5.** Analysis of gene families of different species.

| Species | Genes number | Genes in families | Unclustered genes | Family number | Unique families | Average genes per family |
|---|---|---|---|---|---|---|
| *A. thaliana* | 26,637 | 23,011 | 3626 | 12,620 | 769 | 1.82 |
| *A. altilis* | 33,986 | 27,354 | 6632 | 15,614 | 136 | 1.75 |
| *A. heterophyllus* | 35,845 | 28,969 | 6876 | 15,768 | 242 | 1.84 |
| *F. vesca* | 34,301 | 26,703 | 7598 | 15,188 | 1427 | 1.76 |
| *M. domestica* | 61,721 | 45,647 | 16074 | 17,385 | 3352 | 2.63 |
| *M. notabilis* | 27,085 | 20,805 | 6280 | 14,955 | 567 | 1.39 |
| *P. mume* | 31,128 | 25,702 | 5426 | 16,060 | 566 | 1.6 |
| *P. persica* | 28,701 | 25,385 | 3316 | 15,654 | 231 | 1.62 |
| *Z. jujuba* | 36,942 | 34,050 | 2892 | 14,170 | 755 | 2.4 |

**Table S6.** Enriched GO terms (level 3) of genes in families with expansion.

| **Species** | **GO ID** | **GO Term** | **Type** | **P-value** | **Number of genes** |
|---|---|---|---|---|---|
| | GO:0036094 | small molecule binding | Molecular Function | 5.74E-27 | 862 |
| | GO:0043167 | ion binding | Molecular Function | 4.10E-22 | 1239 |
| | GO:0016740 | transferase activity | Molecular Function | 1.43E-11 | 763 |
| | GO:0097159 | organic cyclic compound binding | Molecular Function | 2.28E-07 | 1274 |
| *A. altilis* | GO:1901363 | heterocyclic compound binding | Molecular Function | 2.28E-07 | 1274 |
| | GO:0001871 | pattern binding | Molecular Function | 0.000227 | 13 |
| | GO:0005515 | protein binding | Molecular Function | 0.003724 | 630 |
| | GO:0016049 | cell growth | Biological Process | 1.21E-06 | 12 |
| | GO:0044700 | single organism signaling | Biological Process | 0.000533 | 72 |

| | | | | | |
|---|---|---|---|---|---|
| | GO:0022857 | transmembrane transporter activity | Molecular Function | 1.45E-17 | 332 |
| | GO:0022892 | substrate-specific transporter activity | Molecular Function | 1.53E-09 | 151 |
| | GO:0016491 | oxidoreductase activity | Molecular Function | 8.81E-07 | 547 |
| | GO:0036094 | small molecule binding | Molecular Function | 2.11E-06 | 983 |
| | GO:0038023 | signaling receptor activity | Molecular Function | 1.00E-05 | 27 |
| | GO:0048037 | cofactor binding | Molecular Function | 0.000102 | 164 |
| | GO:0019208 | phosphatase regulator activity | Molecular Function | 0.000673 | 15 |
| | GO:0016829 | lyase activity | Molecular Function | 0.002478 | 86 |
| | GO:0030246 | carbohydrate binding | Molecular Function | 0.006209 | 63 |
| | GO:0005515 | protein binding | Molecular Function | 0.008019 | 892 |
| | GO:0003682 | chromatin binding | Molecular Function | 0.014389 | 17 |
| | GO:0044703 | multi-organism reproductive process | Biological Process | 1.28E-38 | 96 |
| | GO:0044706 | multi-multicellular organism process | Biological Process | 1.28E-38 | 96 |
| *A. heterophyllus* | GO:0048610 | cellular process involved in reproduction | Molecular Function | 2.68E-37 | 97 |
| | GO:0022414 | reproductive process | Biological Process | 3.57E-36 | 96 |
| | GO:0044707 | single-multicellular organism process | Biological Process | 2.67E-27 | 100 |
| | GO:0044700 | single organism signaling | Biological Process | 2.31E-14 | 149 |
| | GO:0044763 | single-organism cellular process | Biological Process | 3.69E-11 | 831 |
| | GO:0051716 | cellular response to stimulus | Biological Process | 1.07E-08 | 180 |
| | GO:0044765 | single-organism transport | Biological Process | 5.90E-06 | 438 |
| | GO:0044710 | single-organism metabolic process | Biological Process | 2.90E-05 | 739 |
| | GO:0051234 | establishment of localization | Biological Process | 0.000225 | 493 |
| | GO:0009605 | response to external stimulus | Biological Process | 0.002533 | 7 |
| | GO:0051606 | detection of stimulus | Biological Process | 0.007408 | 5 |
| | GO:0031224 | intrinsic to membrane | Cellular Component | 4.87E-06 | 326 |
| | GO:0008287 | protein serine/threonine phosphatase complex | Cellular Component | 0.001066 | 13 |

| | | | | |
|---|---|---|---|---|
| GO:0044425 | membrane part | Cellular Component | 0.003813 | 362 |
| GO:0044421 | extracellular region part | Cellular Component | 0.007408 | 5 |

**Table S7.** Enriched pathways of genes in families with expansion.

| Species | Pathway ID | KEGG description | Number of genes | P-value (<=0.05) |
|---|---|---|---|---|
| *A. altilis* | ko04626 | Plant-pathogen interaction | 265 | 1.33679E-15 |
| | ko04144 | Endocytosis | 127 | 1.55321E-11 |
| | ko04146 | Peroxisome | 79 | 7.69212E-10 |
| | ko04141 | Protein processing in endoplasmic reticulum | 179 | 1.75141E-06 |
| | ko03040 | Spliceosome | 150 | 9.27025E-06 |
| | ko00300 | Lysine biosynthesis | 16 | 6.66998E-05 |
| | ko00450 | Selenocompound metabolism | 22 | 0.000214547 |
| | ko00072 | Synthesis and degradation of ketone bodies | 8 | 0.000814513 |
| | ko00190 | Oxidative phosphorylation | 74 | 0.002151087 |
| | ko00195 | Photosynthesis | 34 | 0.006288728 |
| | ko00260 | Glycine, serine and threonine metabolism | 44 | 0.00738846 |
| | ko00564 | Glycerophospholipid metabolism | 50 | 0.007535005 |
| | ko03060 | Protein export | 28 | 0.007913934 |
| | ko03050 | Proteasome | 30 | 0.008523159 |
| *A. heterophyllus* | ko03430 | Mismatch repair | 88 | 1.09E-14 |
| | ko04626 | Plant-pathogen interaction | 339 | 1.9E-13 |
| | ko03030 | DNA replication | 98 | 6E-13 |
| | ko03440 | Homologous recombination | 84 | 1.28E-12 |
| | ko00380 | Tryptophan metabolism | 40 | 3.05E-08 |
| | ko01110 | Biosynthesis of secondary metabolites | 811 | 1.21E-07 |
| | ko03420 | Nucleotide excision repair | 92 | 1.64E-07 |

| | | | |
|---|---|---|---|
| ko00908 | Zeatin biosynthesis | 59 | 2.94E-06 |
| ko00941 | Flavonoid biosynthesis | 90 | 7.46E-06 |
| ko00052 | Galactose metabolism | 80 | 2.83E-05 |
| ko04712 | Circadian rhythm - plant | 78 | 4.5E-05 |
| ko03040 | Spliceosome | 185 | 5.32E-05 |
| ko00940 | Phenylpropanoid biosynthesis | 189 | 5.41E-05 |
| ko00903 | Limonene and pinene degradation | 36 | 6.01E-05 |
| ko00620 | Pyruvate metabolism | 85 | 7.13E-05 |
| ko00130 | Ubiquinone and other terpenoid-quinone biosynthesis | 53 | 0.000353 |
| ko04144 | Endocytosis | 136 | 0.00072 |
| ko00604 | Glycosphingolipid biosynthesis - ganglio series | 31 | 0.000844 |
| ko02010 | ABC transporters | 74 | 0.000855 |

**Table S8. The gene list of starch biosynthesis in Glycine max.**

| Category | ID in Glycine max |
|---|---|
| AGPL | Glyma.04G011900 |
| AGPL | Glyma.04G030300 |
| AGPL | Glyma.06G011700 |
| AGPL | Glyma.06G030400 |
| AGPL | Glyma.11G116600 |
| AGPL | Glyma.12G042400 |
| AGPL | Glyma.17G252500 |
| AGPL | Glyma.19G223100 |
| AGPS | Glyma.02G304500 |
| AGPS | Glyma.14G009300 |
| BE | Glyma.03G192300 |
| BE | Glyma.04G017700 |
| BE | Glyma.06G018000 |
| BE | Glyma.19G192800 |
| DPE | Glyma.03G121100 |
| DPE | Glyma.04G219500 |
| DPE | Glyma.06G146400 |
| DPE | Glyma.19G125800 |
| GBSS | Glyma.07G049900 |
| GBSS | Glyma.20G218100 |

| | |
|---|---|
| ISA | Glyma.03G151200 |
| ISA | Glyma.06G100600 |
| ISA | Glyma.08G028400 |
| ISA | Glyma.19G153700 |
| PHOH | Glyma.08G334000 |
| PHOH | Glyma.13G057800 |
| PHOH | Glyma.13G235600 |
| PHOH | Glyma.18G067200 |
| PHOH | Glyma.19G028400 |
| PHOH | Glyma.20G026700 |
| PUL | Glyma.10G197000 |
| SS | Glyma.04G235200 |
| SS | Glyma.05G127800 |
| SS | Glyma.06G129400 |
| SS | Glyma.07G260500 |
| SS | Glyma.08G082600 |
| SS | Glyma.13G062700 |
| SS | Glyma.13G204700 |
| SS | Glyma.15G108000 |
| SS | Glyma.19G022900 |
